# Dairy cattle herds mount a characteristic antibody response to highly pathogenic H5N1 avian influenza viruses

**DOI:** 10.1101/2025.04.01.646587

**Authors:** Lindsey R. Robinson-McCarthy, Holly C. Simmons, Aaron L. Graber, Carly N. Marble, Grace W. Graudin, Kevin R. McCarthy

## Abstract

An unprecedented outbreak of a highly pathogenic avian influenza virus, H5 clade 2.3.4.4b, was reported in United States dairy cattle during the spring of 2024. It has now spread to hundreds of herds across multiple states. In humans, antibodies to the hemagglutinin (HA) protein confer the strongest protection against infection. Human herd immunity limits viral spread but also drives the emergence of antigenic variants that escape dominant antibody responses. We used store-bought milk to profile the collective H5N1 antibody response of dairy cattle herds. We detected HA binding antibodies in specific samples from states with recent/ongoing outbreaks. These antibodies present in milk neutralized replicating virus expressing dairy cattle HA and neuraminidase (NA). Despite originating from independent vendors, dairies/plants, geographic regions, and time, antibodies present in these samples are remarkably similar in activity and HA binding specificity. The dominant antibody response was clade 2.3.4.4b HA specific, followed by cross-reactivity with other H5s. Whether the uniformity of the response is a pathway to achieve herd immunity or an avenue for antigenic variants to rapidly escape remains to be seen.

**SIGNIFICANCE:** Establishing human herd immunity ends pandemics. For influenza viruses, this immunity drives continued antigenic evolution that enables viruses to infect once-immune individuals. An outbreak of highly pathogenic avian influenza virus was detected in dairy cattle in 2024 and has spread rapidly across herds and states. We report approaches to assess dairy cattle herd immunity using store-bought milk samples. Across samples separated by geography and time we find dairy cattle mount a strikingly similar antibody response that is strongest to the dairy cattle virus. Benchmarking immunity at this phase of the outbreak is important to understand either eradication or the emergence of antigenic variants that enable reinfection.

## INTRODUCTION

Highly pathogenic avian influenza (HPAI) viruses devastate commercial poultry flocks and can cause high rates of lethal disease in humans. This phenotype is dictated by the hemagglutinin protein (HA), which mediates cell entry (1). Influenza A HAs are classified numerically into 19 subtypes. A second glycoprotein, neuraminidase (NA), facilitates release and shares similar nomenclature. In 2020, an HPAI lineage from H5 clade 2.3.4.4b emerged and is responsible for the current panzootic involving six continents (2). An unprecedented outbreak in United States dairy cattle was detected in the spring of 2024 and has since spread to at least 993 herds in 17 states as of March 26, 2025 (3–5). Serologic data suggest that a far greater number of agricultural/veterinary workers have been infected than the 70+ documented human cases (6–9).

Historically, antigenically novel influenza viruses from animal reservoirs have triggered human pandemics (10, 11). Descendants of these viruses then circulate as endemic, seasonal viruses by acquiring mutations that confer resistance to antibody-mediated herd immunity elicited by previous infection and vaccination (12–16). An understanding of the dairy cattle antibody response to HPAI is needed to forecast the trajectory of the current outbreak and evaluate the consequences of viral antigenic variation. However, access to samples to widely profile immunity of cattle herds is limited.

To overcome these challenges, we developed approaches to profile antibody responses from cattle herds using store-bought milk. We detected H5 antibodies in specific samples from states with known H5N1 dairy cattle infections. Antibodies from milk can block cell entry and inhibit viral spread of a replicating virus expressing dairy cattle H5N1 HA and NA. We find that across samples from independent vendors, brands, dairies/plants, geographic regions and time, antibodies present in these samples are remarkably similar in their pattern of HA reactivity.

Clade 2.3.4.4b specific antibodies dominate the response, with some cross-reactivity to other divergent H5s and less to HAs from different subtypes. We conclude that at the herd level, exposure to clade 2.3.4.4b viruses elicits this strikingly similar, stereotypic, antibody response. Whether this immunity will be sufficient to protect against current and future H5 viruses or drive the emergence of antigenic variants that escape it warrants increased surveillance.

## RESULTS

### Identification of H5-reactive antibodies in store-bought milk

Reports indicate that dairy cattle H5N1 viruses rapidly spread through herds (3, 4, 17). Following infection, antibodies to this virus should be present in the milk of lactating animals (18, 19). Given the number of cows likely exposed to H5N1 in affected herds (4, 5, 17, 20), we reasoned that H5 specific antibodies could be detected in store-bought milk. We obtained seven samples from California and two from Colorado in early October 2024 (Milk 1 to Milk 7) (Table 1). These were chosen to encompass multiple milk fat percentages, dairies/processing plants, and expiration dates (as a proxy for processing dates). Milk samples were tested for reactivity with a recombinantly expressed, soluble, HA ectodomain trimer derived from A/dairy cow/Texas/24-008749-001/2024 (H5N1) in enzyme-linked immunosorbent assays (ELISA). We used milk from Pennsylvania, where no cases had been reported at the time of purchase, as a negative binding control and an H5-binding human monoclonal antibody FLD194 (21) as a positive binding control. One Colorado and one California milk sample demonstrated concentration-dependent H5 binding, which we defined as H5 antibody positive (Table 1, Figure S1A).

**Table 1.**
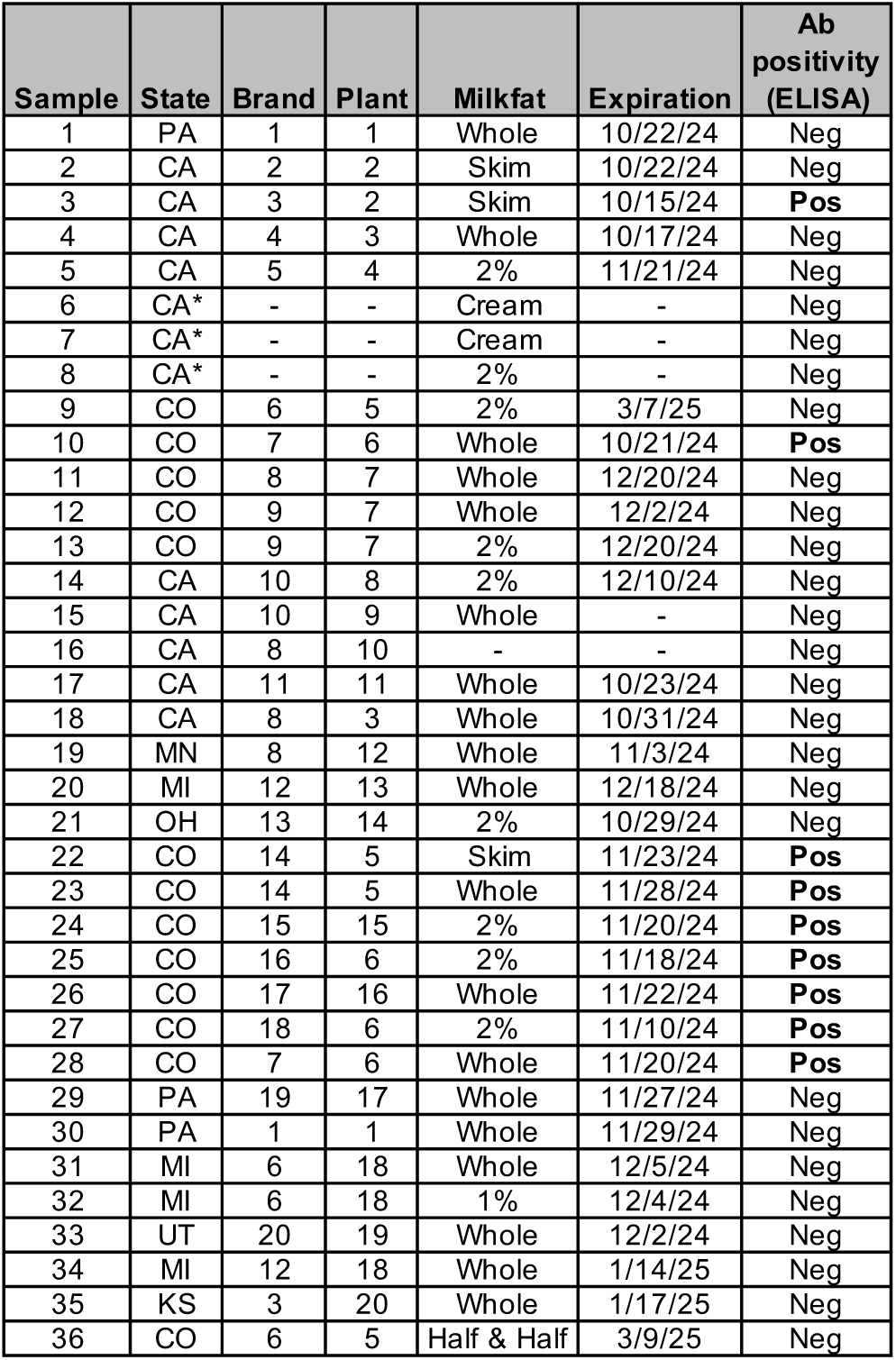
Store-bought milk samples contain H5 HA-reactive antibodies. Milk samples were collected from multiple states. “-”: information was unavailable. “*” Samples 6-8 are presumed to be from California, but brand and processing plant information were not available. Antibody positivity was determined by ELISA with A/dairy cow/Texas/24-008749-001/2024 H5 HA (See Figure S1).

| Sample | State | Brand | Plant | Milkfat | Expiration | Ab positivity (ELISA) |
| --- | --- | --- | --- | --- | --- | --- |
| 1 | PA | 1 | 1 | Whole | 10/22/24 | Neg |
| 2 | CA | 2 | 2 | Skim | 10/22/24 | Neg |
| 3 | CA | 3 | 2 | Skim | 10/15/24 | Pos |
| 4 | CA | 4 | 3 | Whole | 10/17/24 | Neg |
| 5 | CA | 5 | 4 | 2% | 11/21/24 | Neg |
| 6 | CA* | - | - | Cream | - | Neg |
| 7 | CA* | - | - | Cream | - | Neg |
| 8 | CA* | - | - | 2% | - | Neg |
| 9 | CO | 6 | 5 | 2% | 3/7/25 | Neg |
| 10 | CO | 7 | 6 | Whole | 10/21/24 | Pos |
| 11 | CO | 8 | 7 | Whole | 12/20/24 | Neg |
| 12 | CO | 9 | 7 | Whole | 12/2/24 | Neg |
| 13 | CO | 9 | 7 | 2% | 12/20/24 | Neg |
| 14 | CA | 10 | 8 | 2% | 12/10/24 | Neg |
| 15 | CA | 10 | 9 | Whole | - | Neg |
| 16 | CA | 8 | 10 | - | - | Neg |
| 17 | CA | 11 | 11 | Whole | 10/23/24 | Neg |
| 18 | CA | 8 | 3 | Whole | 10/31/24 | Neg |
| 19 | MN | 8 | 12 | Whole | 11/3/24 | Neg |
| 20 | MI | 12 | 13 | Whole | 12/18/24 | Neg |
| 21 | OH | 13 | 14 | 2% | 10/29/24 | Neg |
| 22 | CO | 14 | 5 | Skim | 11/23/24 | Pos |
| 23 | CO | 14 | 5 | Whole | 11/28/24 | Pos |
| 24 | CO | 15 | 15 | 2% | 11/20/24 | Pos |
| 25 | CO | 16 | 6 | 2% | 11/18/24 | Pos |
| 26 | CO | 17 | 16 | Whole | 11/22/24 | Pos |
| 27 | CO | 18 | 6 | 2% | 11/10/24 | Pos |
| 28 | CO | 7 | 6 | Whole | 11/20/24 | Pos |
| 29 | PA | 19 | 17 | Whole | 11/27/24 | Neg |
| 30 | PA | 1 | 1 | Whole | 11/29/24 | Neg |
| 31 | MI | 6 | 18 | Whole | 12/5/24 | Neg |
| 32 | MI | 6 | 18 | 1% | 12/4/24 | Neg |
| 33 | UT | 20 | 19 | Whole | 12/2/24 | Neg |
| 34 | MI | 12 | 18 | Whole | 1/14/25 | Neg |
| 35 | KS | 3 | 20 | Whole | 1/17/25 | Neg |
| 36 | CO | 6 | 5 | Half & Half | 3/9/25 | Neg |

To identify further H5 antibody positive samples, we purchased 24 additional milk products from states with reported dairy cattle H5N1 infections. This included milk from 8 states bottled in 20 dairy/milk plants. A focused effort was made to acquire additional products from Colorado and California. Additional Pennsylvania negative controls were also obtained. Only seven samples, all from Colorado, had detectable levels of H5 HA reactivity (Table 1, Figure S1B). Notably, all of the Colorado products with undetectable H5 binding antibodies were ultra-pasteurized while those with detectable binding were not (Table S1). None of the additional California milk samples had detectable H5 HA binding antibodies. The one positive sample from California was collected before the scale of the outbreak was recognized and hundreds of herds were quarantined (22).

### Antibodies in milk have antiviral activity

We used a replicating recombinant vesicular stomatitis virus (rVSV-H5N1dc2024) that expresses the H5 and N1 from A/dairy cow/Texas/24-008749-001/2024 in place of its glycoprotein (23) to assess the antiviral activity of antibodies in milk. This virus also expresses a green fluorescent protein (GFP) reporter to track infection. We performed neutralization assays using the nine H5 antibody-positive milk samples and two Pennsylvania negative controls (Figure 1A and S2). In these assays, the inoculum and milk remained on cells for the duration of the experiment. Milk concentrations did not exceed 10% v/v of the cell culture media. Only one, sample 10 from Colorado, completely inhibited infection at dilutions of 1:10 and 1:20, although most had measurable half-maximal inhibitory concentrations (IC50). However, for all H5 antibody-positive samples, we observed a concentration-dependent reduction the number of infected foci and cell-cell spread over the course of two days (Figure 1A-B). Antibodies in milk likely inhibit viral replication through multiple mechanisms, including directly blocking cell entry as seen by a reduction in the number of GFP foci, and by interfering with assembly or release of progeny virions as seen by the reduction in spread. In store-bought milk, which is pooled from potentially naive animals or ones in various states of convalescence, the concentrations of inhibitory antibodies appear relatively low.

**Figure 1.**
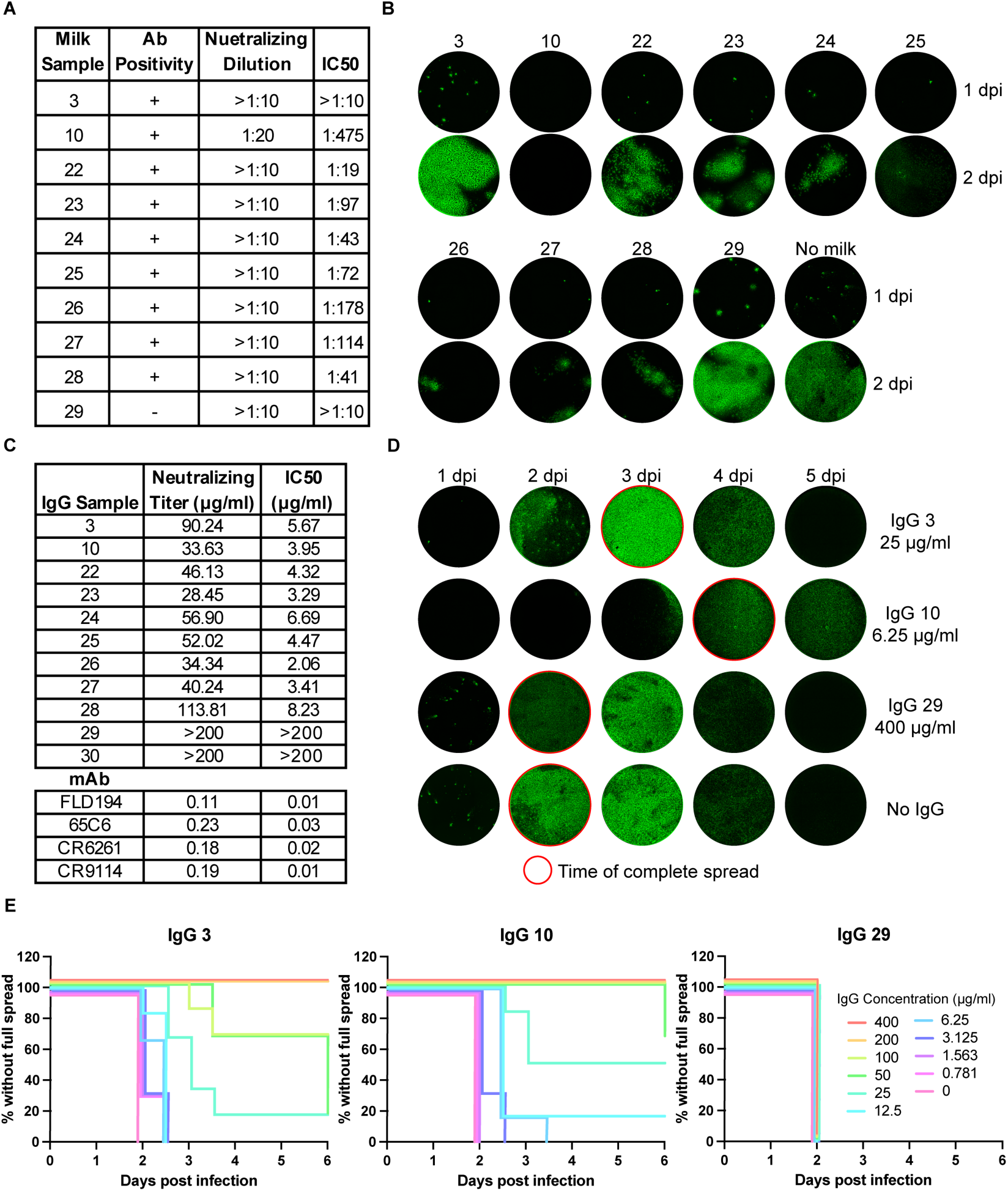
Antibodies in milk have antiviral activity. (A) Dilution at which each milk sample (Table 1) completely inhibited infection by rVSV-H5N1dc2024, and IC50 neutralizing titers for each milk sample. (B) Representative images of rVSV-H5N1dc2024 infected GFP-expressing cells at 1- and 2-days post infection in the presence of each milk sample at a 1:10 dilution. (C) Neutralizing titers of purified IgG and mAbs (21, 33–35) for rVSV-H5N1dc2024. IgG sample numbers match the milk sample from which they were purified. (D) Representative images of rVSV-H5N1dc2024 infected cells in the presence of sub-neutralizing concentrations of purified IgG over 5 days. IgG concentration for each is indicated. Wells that have reached complete viral spread are outlined in red. Images taken at half-day intervals are not shown. (E) Protection from viral spread by purified milk IgG. Spread of rVSV-H5N1dc2024 infection in the presence of purified IgG was assessed twice per day over 5 days for each concentration of IgG. “Survival” was defined as conditions where infection had not spread throughout the well.

We purified and concentrated IgG from these milk samples to further characterize their inhibitory functions. At the highest concentrations, antibodies from all nine H5 antibody-positive samples fully inhibited infection, while those from negative samples had no antiviral activity (Figure 1C). We determined the minimal antibody concentration required to completely block infection and IC50 values (Figure S2). Among our samples, these titers varied over a four-fold range. We then assessed the time needed for viruses to spread across the entire cell monolayer of an infected well. Inhibition of cell-cell spread occurred at greater dilutions than those required to fully prevent infection (Figure 1D-E and Figure S3).

### Dairy cattle mount a common antibody response to H5N1 viruses

We profiled the HA-reactivity of antibodies in milk using a panel of 83 recombinantly expressed, soluble HA ectodomain trimers (Table S2). These include the matched A/dairy cow/Texas/24-008749-001/2024 HA, a second clade 2.3.4.4b H5 HA, increasingly divergent H5 HAs from different clades/lineages, endemic human and animal H2, H1, H3, avian-origin H7, and human influenza B HAs. All nine ELISA-positive milk samples reacted most strongly with clade 2.3.4.4b HAs (Figure S4). A minimal, non-specific, “background” level of HA reactivity was observed in our negative control samples.

While the magnitude of reactivity to each HA differed across samples, we observed a strikingly consistent hierarchy of antibody responses across samples, time, plant/dairy, and states (Figure 2). Binding was strongest to clade 2.3.4.4b HAs, followed by other H5s, H2s, H1s. There was minimal reactivity with group 2 H3 and H7 HAs. The H5 specificity was more pronounced with milk as the analyte than with purified IgG, although both were strongly biased to group 1 HAs (Figures S4 and S5).

**Figure 2.**
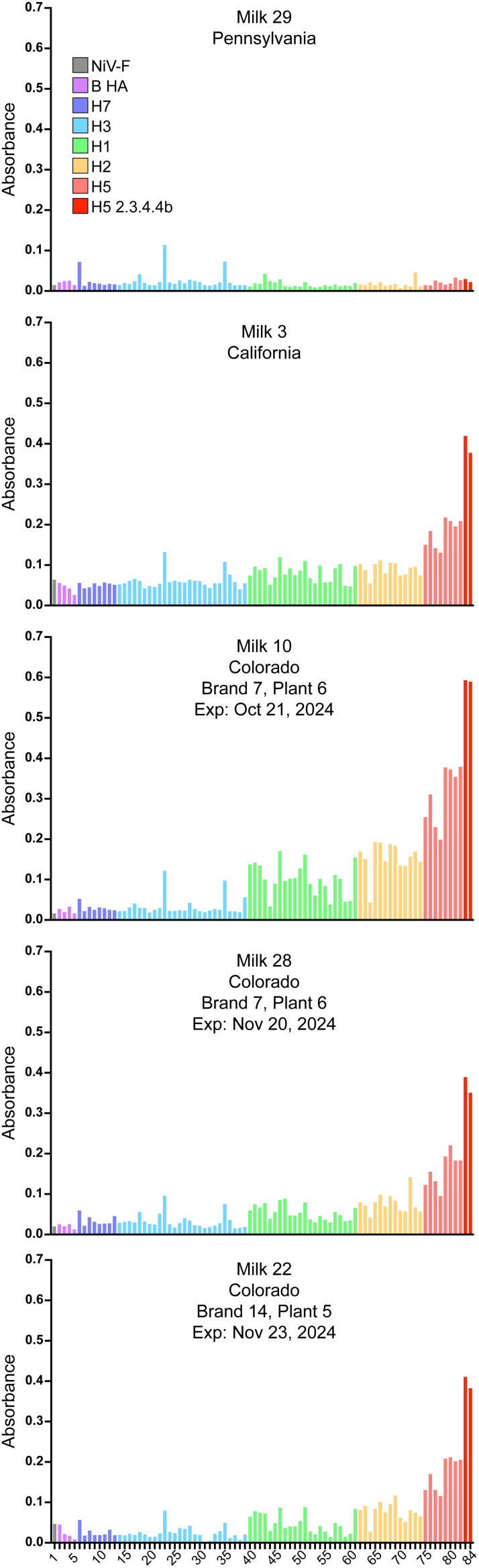
Breadth of HA reactivity of milk antibodies. Binding of antibodies in milk to recombinant full-length secreted ectodomain HA trimers from 83 unique isolates was assessed by ELISA. Nipah virus F protein (NiV-F) was included as a negative control. Bound antibody was detected using HRP-conjugated protein A/G. Bars are colored based on HA subtype, with H5 clade 2.3.4.4.b H5 in darker red for emphasis. The identity of each HA is provided in Table S2.

We performed ELISA titrations with the purified IgGs because they could be used at higher concentrations and normalized for protein content (Figures S6 and S7). Half maximal effective concentrations (EC_50_) were lowest for clade 2.3.4.4b HAs (Figure 3). EC_50_ values were approximately 2-fold higher for other H5s and nearly 5-fold higher for H1s and H2s. Reactivity with H3 HA was minimal and similar to the reactivity of IgGs purified from the negative control Pennsylvania milk. Together, these data suggest that the strongest antibody response elicited by infection is H5 clade 2.3.4.4b-dominant, sensitive to variation within H5 HAs, and strongly group 1 HA biased.

**Figure 3.**
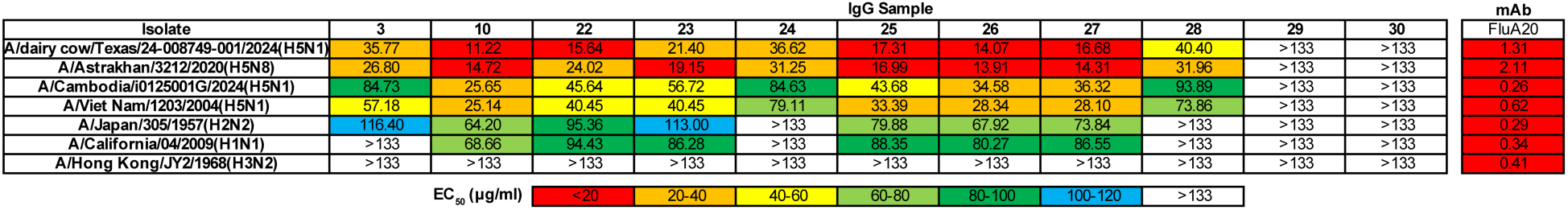
Purified IgG from milk reacts most strongly to clade 2.3.4.4.b H5 HAs. ELISA titrations were performed to determine the EC50 values for each purified IgG sample to H5, H2, H1, and H3 HAs. A/dairy cow/Texas/24-008749-001/2024 and A/Astrakhan/3212/2020 both belong to clade 2.3.4.4b. mAb FluA-20 (36) was included as a broadly binding control antibody.

## DISCUSSION

An outbreak of HPAI in dairy cattle is unprecedented. The paucity of samples from confirmed H5N1 convalescent dairy cows has hindered the characterization of their antibody response to H5 clade 2.3.4.4b viruses. To overcome this challenge, we developed methods to profile the H5N1 antibody response at the scale of herd(s). Using store-bought milk, we created a snapshot of dairy cattle herd immunity elicited by a primary H5N1 exposure. Our “immunosurveillance” approach is generalizable and well-suited for tracking the trajectory of exposure and immunity in this H5N1 outbreak or other endemic and emerging pathogens over time. Our ELISA based assays and use of rVSV-H5N1dc2024 enable similar efforts to be performed in most molecular biology laboratories with limited need for specialized equipment or high biocontainment facilities.

Antibodies present in all of our H5 antibody-positive milk samples inhibit viral replication through multiple mechanisms, but were present at relatively low concentrations. Milk from experimentally infected cows had maximal neutralizing titers between 8 and 25-fold higher than our most neutralizing bulk milk sample (18, 19). Using the GFP reporter present in VSV-H5N1dc2024 we detected and quantified the activity of antibodies that prevent cell entry (neutralizing) and those that prevent cell-cell spread. Antibodies preventing cell-cell spread were inhibitory at sub-neutralizing milk/IgG concentrations. Given the high titers of infectious virus in the milk of actively infected cows prior to pasteurization (4, 18, 19), antibody concentrations in bulk milk tanks may be insufficient to fully neutralize H5N1 viruses and may therefore harbor infectious virus.

We found that the pattern of HA reactivity was remarkably similar across store-bought milk samples from multiple states, processing plants, brands, and time (same brand and plant). This pattern is defined by a strong clade 2.3.4.4b response, diminished binding to other H5s, and weaker binding to H2s and H1s (group 1 HAs). Binding to H3 or H7 (group 2 HAs) was limited or absent. Despite using a large array of recombinant HA proteins that include human and animal isolates, we did not find evidence of unreported, “cryptic”, dairy cattle infections from either H5 or other subtypes tested. Prior to this outbreak, infection of dairy cattle with influenza A viruses appears to be rare as no recall response was detected. Recent reports of dairy cattle infection with genetically distinct clade 2.3.4.4b viruses suggest that these viruses may be particularly well-suited to infecting cows (24–26).

At the herd level, exposure to H5N1 reproducibly elicits a strikingly similar antibody response. Infection, or possibly vaccination with matched strains, are likely to elicit comparable responses in currently naive animals. Given the similarity of antibody responses across distinct herds and the relative magnitude of the H5 clade 2.3.4.4b-specific response, monitoring for antigenic variants that escape herd immunity is warranted.

## METHODS

### Milk samples

Commercially available, store-bought milk was obtained from grocery stores, convenience stores, and catered refreshments. The provenance of the milk was determined by the processing plant code. In instances where no plant code was provided, the dairy was confirmed to operate within the state of purchase. Milk brand and processing plant information was deidentified to adhere to the current Centers of Excellence for Influenza Research and Response (CEIRR) Best Practice Guidelines for the Reporting of Milk Testing and Results (27). Samples 6-8 were obtained from the coffee station at the West Coast Retrovirus Meeting in Palm Springs, California in early October 2024. These samples were assumed to originate in California, but further information about their provenance was not available. Full information for milk samples tested is provided in Table S1.

### Cells and viruses

BSRT7 cells (28) were maintained at 37°C and 5% CO_2_ in Dulbecco’s modified Eagle medium (DMEM; ThermoFisher) supplemented with 10% fetal bovine serum (FBS) and penicillin/streptomycin (pen/strep; ThermoFisher). 293F cells were maintained at 37°C with 8% CO_2_ in FreeStyle 293 Expression Medium (ThermoFisher) supplemented with pen/strep.

rVSV-H5N1dc2024 was previously generated (23) and propagated on BSRT7 cells in viral growth medium (DMEM supplemented with 2% FBS, pen/strep, and 25mM HEPES). rVSV-H5N1dc2024 was titered by plaque assay on BSRT7 cells as previously described (23).

### Plasmids

Synthetic DNA corresponding to the full-length ectodomain (FLsE) of influenza HAs were cloned into a pVRC vector modified to encode a C-terminal thrombin cleavage site, a T4 fibritin (foldon) trimerization tag, and a 6xHis tag (29, 30). Point mutations to stabilize HA as a prefusion trimer were added to specific sequences (annotated in Table S2), based on previously reported stabilizing mutations (31). Synthetic DNA corresponding to a prefusion stabilized Nipah virus F protein (NiV-F) (32) was cloned into pVRC as above. Accession numbers for all HA sequences are provided in Table S2.

Synthetic DNA corresponding to the heavy and light chain variable domains of antibodies FLD194 (21), 65C6 (33), CR9114 (34), CR6261 (35), and FluA-20 (36) were ordered from IDT and cloned into modified pVRC8400 plasmids (29) containing full length human IgG1 heavy chains or human kappa or lambda light chains.

All plasmid sequences were verified through Sanger sequencing (Azenta) or whole plasmid nanopore sequencing (Plasmidsaurus).

### Monoclonal Antibody Production

Recombinant mAbs were produced as previously described (37) by transient transfection of heavy and light chain plasmids into 293F cells using PEI transfection reagent. Five days post-transfection, supernatants were collected, clarified by low-speed centrifugation, and incubated overnight with Protein A Agarose Resin (GoldBio) at 4°C. Resin was collected in a chromatography column and washed with one column volume of 10 mM tris(hydroxymethyl)aminomethane (tris), 150 mM NaCl at pH 7.5. mAbs were eluted in 0.1M Glycine (pH 2.5) which was immediately neutralized by 1M tris (pH 8.5). Antibodies were dialyzed against phosphate buffered saline (PBS) pH 7.4.

### Recombinant HA expression and purification

Recombinant HA FLsE and NiV-F were expressed and purified as previously described (37). Briefly, 293F cells were transiently transfected with FLsE encoding plasmids using PEI. Transfection complexes were prepared in Opti-MEM and added to cells. Supernatants were harvested 4 to 5 days post transfection and clarified by low-speed centrifugation. HA and NiV-F trimers were purified by passage over Co-NTA agarose (Clontech) followed by gel filtration chromatography on Superdex 200 (GE Healthcare) in 10 mM Tris-HCl, 150 mM NaCl at pH 7.5.

### H5 reactive antibody screening

500 ng of recombinant HA FLsE per-well were adhered to 96-well plates (Corning) in PBS pH 7.4 overnight at 4°C. HA-bound plates were washed with 0.05% TWEEN-20 in PBS (PBS-T) and blocked at room temperature for one hour in PBS-T with 2% BSA. Blocking solution was removed and 2-fold dilutions of milk in blocking solution were added to the wells. mAb FLD194 (21), diluted in 5-fold dilutions starting at 500 ng/μl was included on each plate as a positive binding control. Plates were then incubated for one hour at room temperature and then washed four times with PBS-T to remove milk/mAb. 100 μl of blocking buffer with peroxidase conjugated recombinant protein A/G (ThermoFisher) diluted 1:10,000 was added to each well and incubated for one hour at room temperature. Plates were washed three times with PBS-T and developed using 150 μl 1-Step ABTS substrate (ThermoFisher). Following incubation at room temperature, HRP reactions were stopped by the addition of 100 μl of 1% sodium dodecyl sulfate (SDS) solution. Plates were read on a Molecular Devices SpectraMax 340PC384 Microplate Reader at 405 nm. All measurements were performed in single replicates. Background subtracted data were grafted using Prism 10 software (Graphpad).

### Antibody purification from milk

Milk or milk diluted one to one in 10 mM Tris-HCl, 150 mM NaCl at pH 7.5 was incubated with immobilized Protein G resin (Thermo Scientific) overnight at 4°C. The resin was collected in a chromatography column and washed with at least two column volumes of 10 mM Tris-HCl, 150 mM NaCl at pH 7.5. Antibodies were eluted in 0.1M Glycine (pH 2.5), which was immediately neutralized by 1M tris (pH 8.5). Antibodies were then concentrated and dialyzed against phosphate buffered saline (PBS) pH 7.4.

### Milk neutralization assays

Neutralization assays were performed as previously reported, with some modifications (23). Milk samples were diluted in 2-fold serial dilutions in viral growth medium. 100 plaque forming units (pfu) of rVSV-H5N1dc2024 per well was added to diluted milk and samples were incubated for one hour at room temperature. Milk:virus mixture was added to BSRT7 cells, which were incubated at 37°C. Milk:virus mixture was left on cells for the duration of the experiment. Cells were imaged for GFP expression 1 and 2 days post infection using an EVOS automated fluorescence microscope (ThermoFisher) with a 4x objective. Images were analyzed using Image J software (NIH). Neutralizing titers are the dilution of milk that completely inhibited infection by 100 pfu of rVSV-H5N1dc2024. Fluorescent foci per well were counted from images taken one day post-transfection. Data were analyzed and half maximal inhibitory concentration (IC50) values calculated using Prism 10 software (Graphpad).

### Purified IgG neutralization assays

Recombinant monoclonal antibodies and purified IgG from milk were diluted in serial two-fold dilutions and neutralization experiment was performed as described above. Antibody:virus mixture was maintained on the cells throughout the experiment. Cells were imaged twice per day over 5 days. Endpoint neutralizing titers are the concentration of IgG or mAb required to completely inhibit infection by 100 pfu of virus. Fluorescent foci per well were counted from images taken one day post-transfection. Data were plotted and analyzed using Prism 10 software (Graphpad) to determine half maximal inhibitory concentration (IC50). Time to spread was determined for each well as the time point at which the virus had overtaken the entire well. Data were plotted as survival curves using Prism 10 software (Graphpad).

### Antibody binding breadth assay

100 ng of recombinant HA FLsE or NiV-F were adhered in duplicate to 96-well plates (Corning) in PBS pH 7.4 overnight at 4°C. HA bound plates were washed with 0.05% TWEEN-20 in PBS (PBS-T) and blocked at room temperature for one hour in PBS-T with 2% bovine serum albumin (BSA). Blocking solution was removed and either diluted commercial milk samples or purified IgG from milk samples were added to each plate. Milk samples were diluted to 10% in blocking solution and purified IgG was diluted to 1 mg/ml. Diluted milk or IgG was removed after 1 hour. Plates were washed three times with PBS-T and incubated with 1:10,000 peroxidase conjugated recombinant protein A/G (ThermoFisher) in blocking solution for one hour. Following secondary incubation, plates were washed three times with PBS-T and developed with 100 μl of 1-Step Slow TMB-ELISA Substrate Solution (ThermoFisher) for 8 minutes, after which the reaction was stopped with 100 μl of 2M sulfuric acid. Absorbance values were measured using Molecular Devices SpectraMax 340PC384 Microplate Reader at 450 nm.

### Milk derived IgG binding curve ELISAs

500 ng of recombinant HA FLsE were adhered to 96-well plates (Corning) in PBS pH 7.4 overnight at 4°C. HA-bound plates were washed with 0.05% TWEEN-20 in PBS (PBS-T) and blocked at room temperature for one hour in PBS-T with 2% BSA. Blocking solution was removed and 5-fold dilutions of purified IgG from milk were added to the wells. mAb FluA-20 was included on each plate as a positive control and to allow for normalization. Primary antibody was incubated for one hour then removed. Plates were washed three times with PBS-T and incubated with 1:10,000 peroxidase conjugated recombinant protein A/G (ThermoFisher) in blocking solution for one hour. Following secondary incubation, plates were washed three times with PBS-T and developed with 100 μl of 1-Step Slow TMB-ELISA Substrate Solution (ThermoFisher) for 8 minutes, after which the reaction was stopped with 100 μl of 2M sulfuric acid. Absorbance values were measured using Molecular Devices SpectraMax 340PC384 Microplate Reader at 450 nm. All measurements were performed in technical triplicate. Data were plotted and analyzed using Prism 10 software (Graphpad) to determine half maximal effective concentration (EC_50_).

## ACKNOWLEDGEMENTS

We thank Caitlin M. Weaver, Sandra Bos, Alex J. Guseman, and Melissa E. Kane for their help in obtaining samples.

This work was funded by the National Institute of Allergy and Infectious Diseases, Centers of Excellence for Influenza Research and Response, contract number 75N93021C0001 (Option 13F) (to KRM), by National Institute of General Medical Sciences grant 1R35GM154844 (to KRM) and by NIH award (UC7AI180311) from the National Institute of Allergy and Infectious Diseases (NIAID) supporting the Operations of The University of Pittsburgh Regional Biocontainment Laboratory (RBL) within the Center for Vaccine Research (CVR).

---

**Figure S1.**
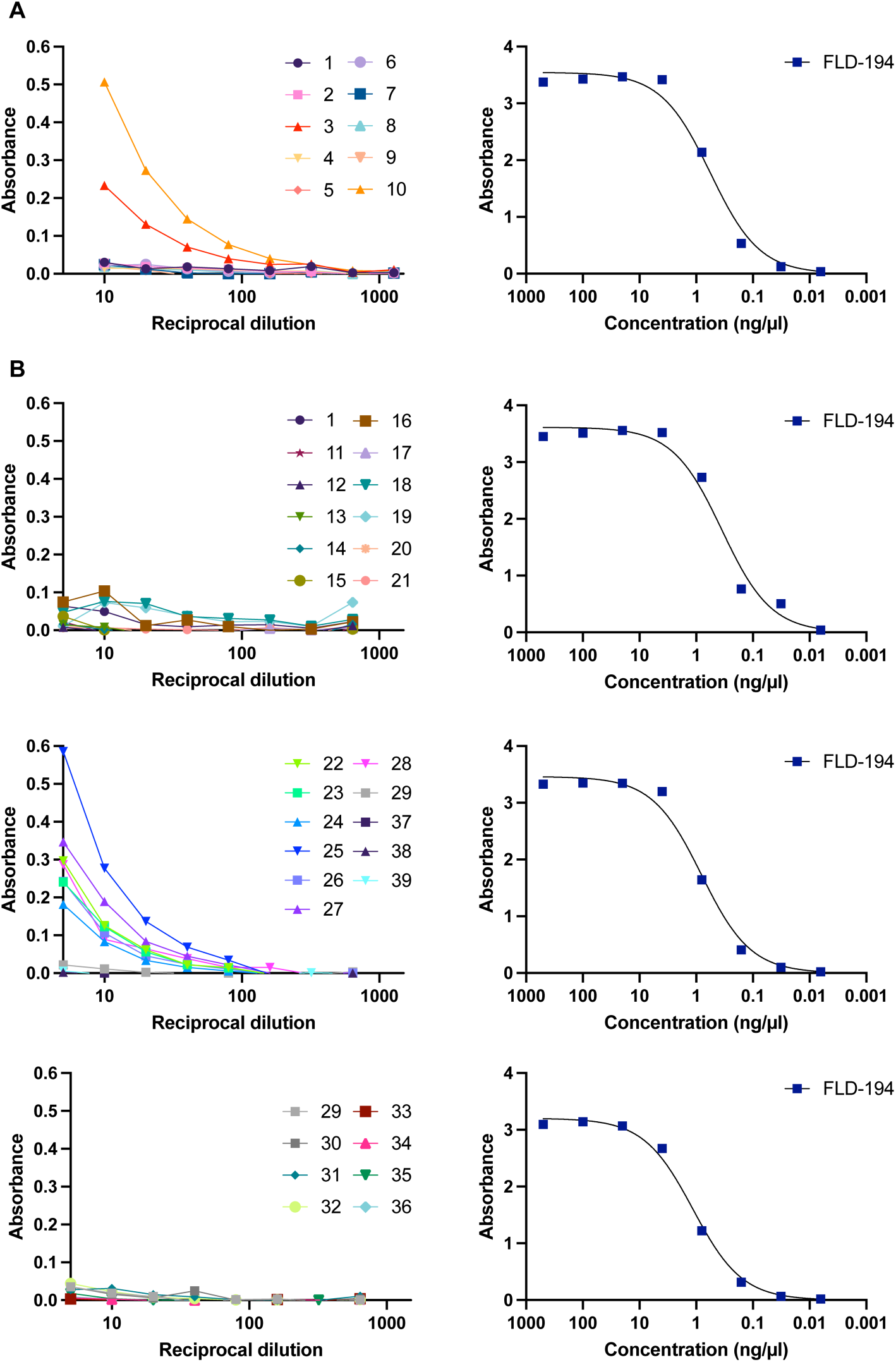
ELISA screening for H5-reactive antibodies in milk. Milk samples were screened over a range of dilutions for binding to A/dairy cow/Texas/24-008749-001/2024 H5 HA. Background subtracted absorbance values are shown for each plate, alongside standard curves for mAb FLD194 (21) for the same plate. (A) Initial California, Colorado, and Pennsylvania samples screened. (B) Additional samples from multiple states.

**Figure S2.**
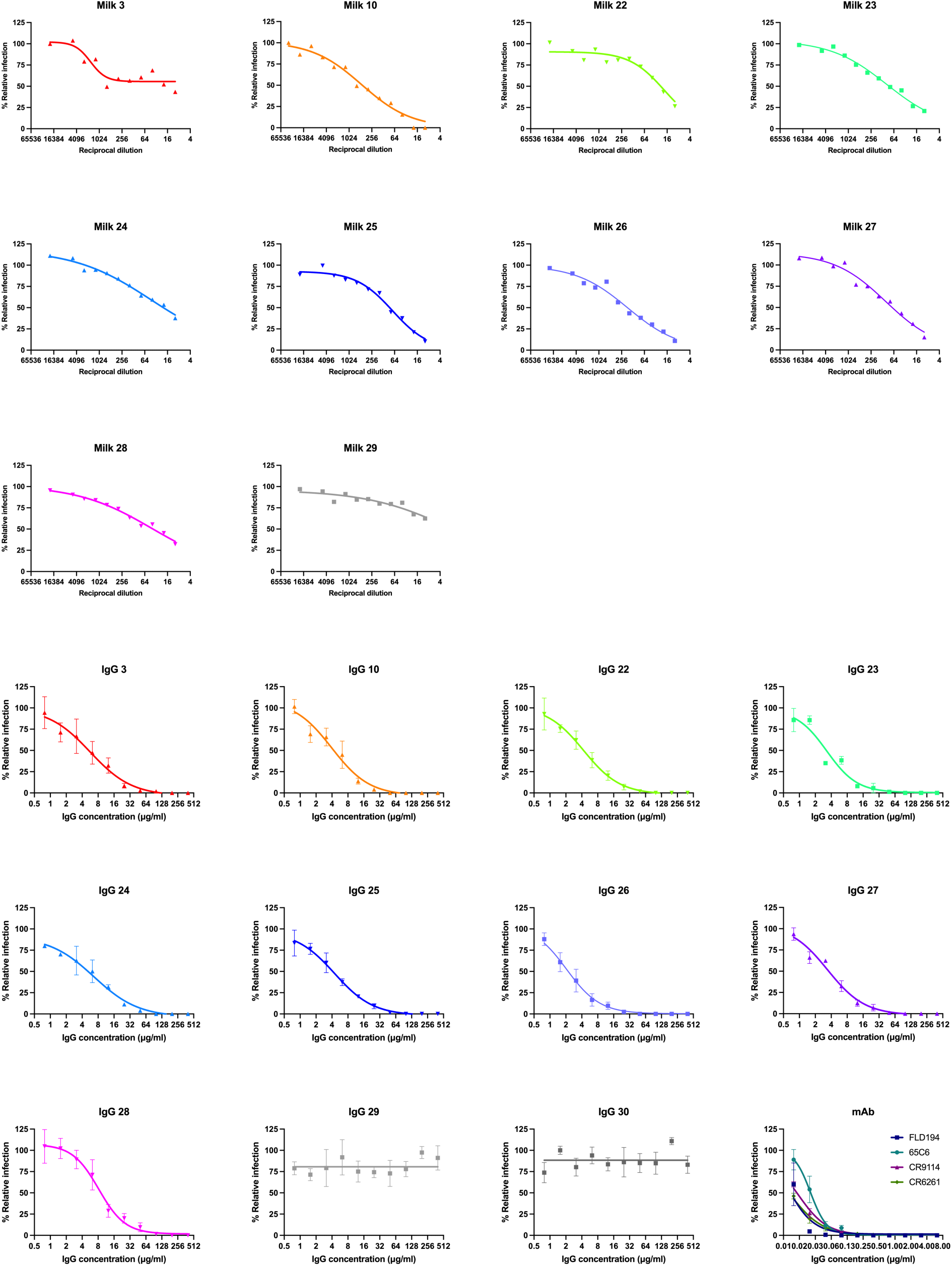
Neutralization of rVSV-H5N1dc2024 by milk and purified milk IgG. Fluorescent focus forming neutralization test was performed. Cells infected with rVSV-H5N1dc2024 in the presence of each concentration of milk, purified IgG, or mAbs (21, 33–35) were imaged and fluorescent foci per well were quantified and normalized to wells with no milk/IgG/mAb to determine percent relative infection. Error bars represent standard error of the mean from three independent experiments. Experiments with milk are averages of two independent experiments.

**Figure S3.**
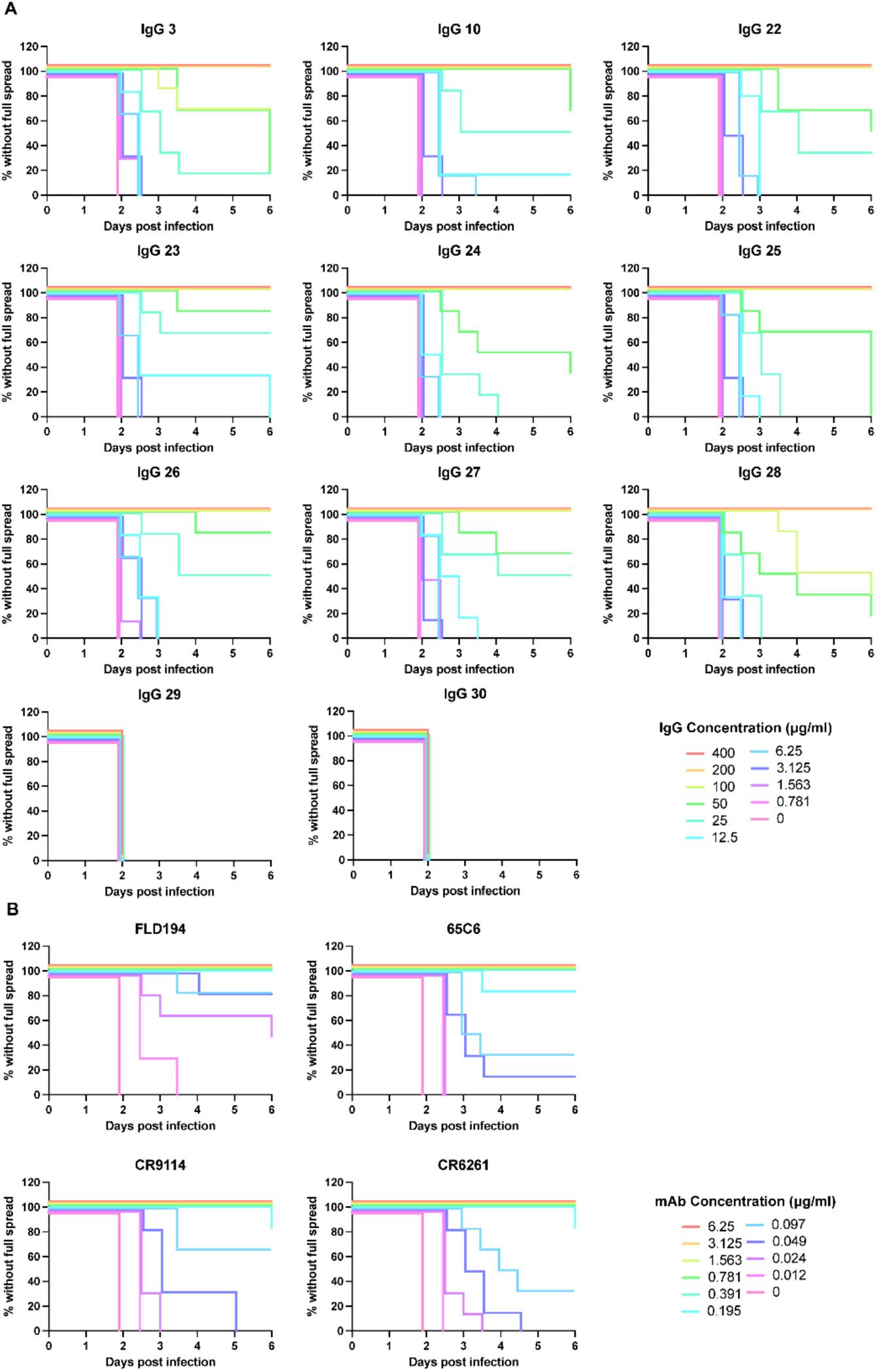
Purified IgG from milk inhibits viral spread at sub-neutralizing concentrations. (A) Spread of rVSV-H5N1dc2024 infection in the presence of purified milk IgG for all samples was assessed twice per day over 5 days for each concentration of IgG. Survival was defined as conditions where infection had not spread throughout the well. Wells which had spreading infection, but which had not reached complete spread by 5 days were annotated as reaching full spread at 6 days. (B) The same assay was performed using H5 HA-reactive mAbs FLD194 (21) (HA head binding), 65C6 (33) (HA head binding), CR9114 (34) (HA stem binding), and CR6261 (35) (HA stem binding). Each group represent N=6 total measurements from three independent experiments.

**Figure S4.**
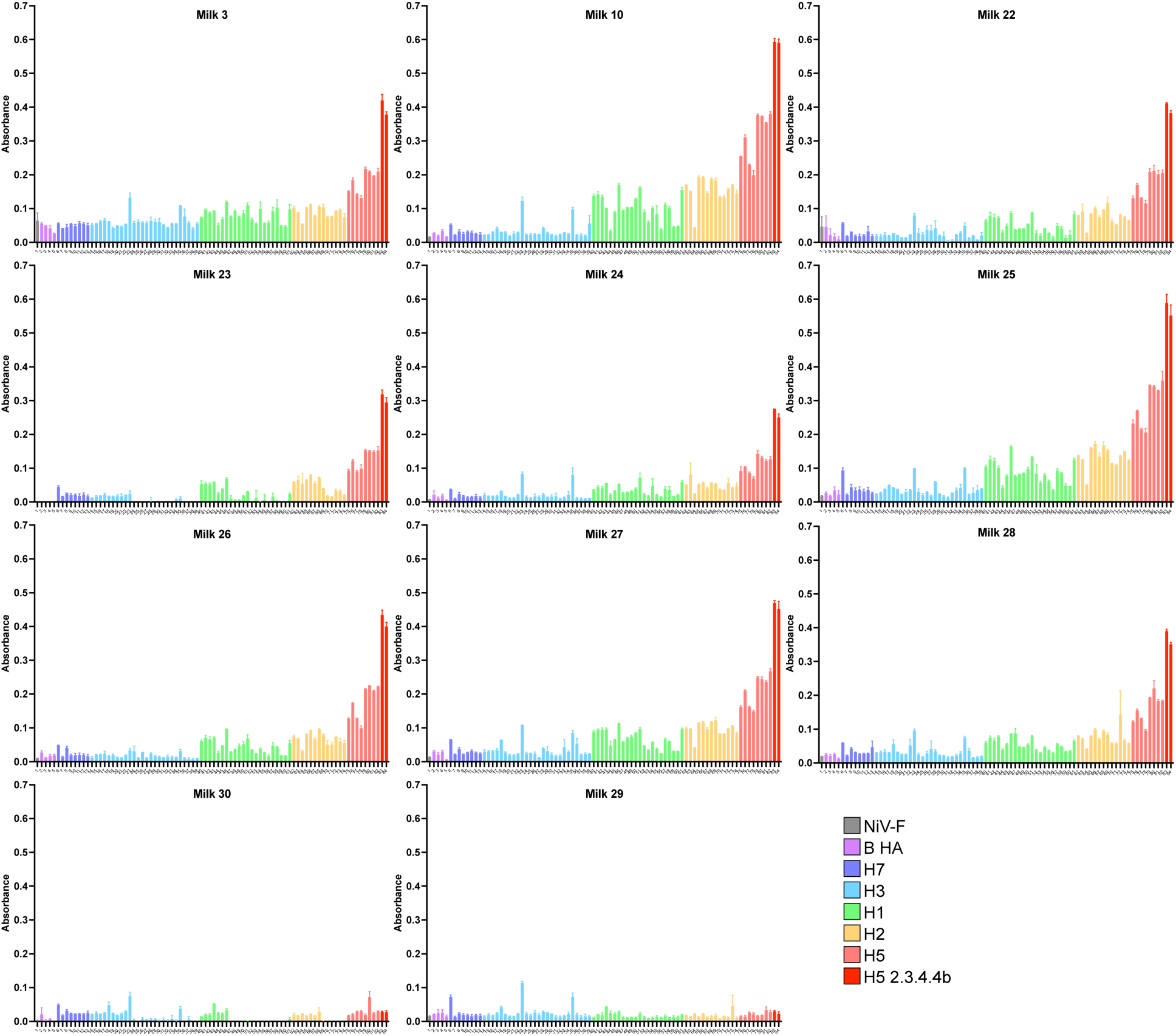
HA reactivity breadth of milk antibodies. Reactivity of milk antibodies to recombinant full-length secreted ectodomain HA trimers from 83 unique isolates was assessed by ELISA. Nipah virus F protein (NiV-F) was included as a negative control. Bound antibody was detected using HRP-conjugated protein A/G. Bars are colored based on HA subtype, with H5 clade 2.3.4.4.b H5 in darker red for emphasis. The identity of each HA is provided in Table S2.

**Figure S5.**
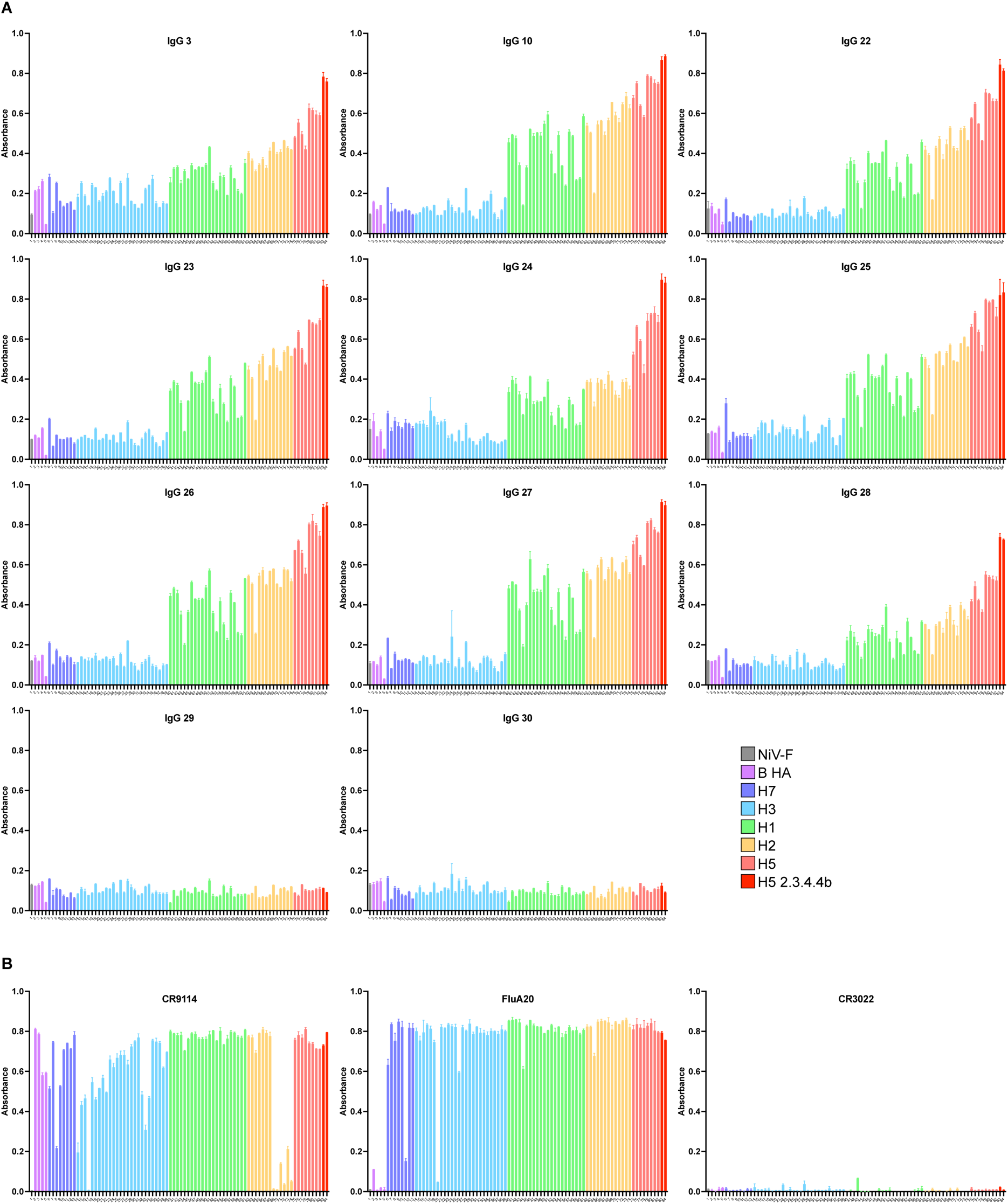
Breadth of HA reactivity of purified IgG from milk samples. (A) Reactivity of purified IgG from milk to recombinant full-length secreted ectodomain HA trimers from 83 unique isolates was assessed by ELISA. Nipah virus F protein (NiV-F) was included as a negative control. Bound antibody was detected using HRP-conjugated protein A/G. Bars are colored based on HA subtype, with H5 clade 2.3.4.4.b H5 in darker red for emphasis. The identity of each HA is provided in Table S2. (B) Binding of mAbs CR9114 (34) (broad HA stem binding), FluA-20 (36) (broad HA head binding), and CR3022 (38) (SARS-CoV spike binding) to validate HAs tested using the same ELISA as in (A).

**Figure S6.**
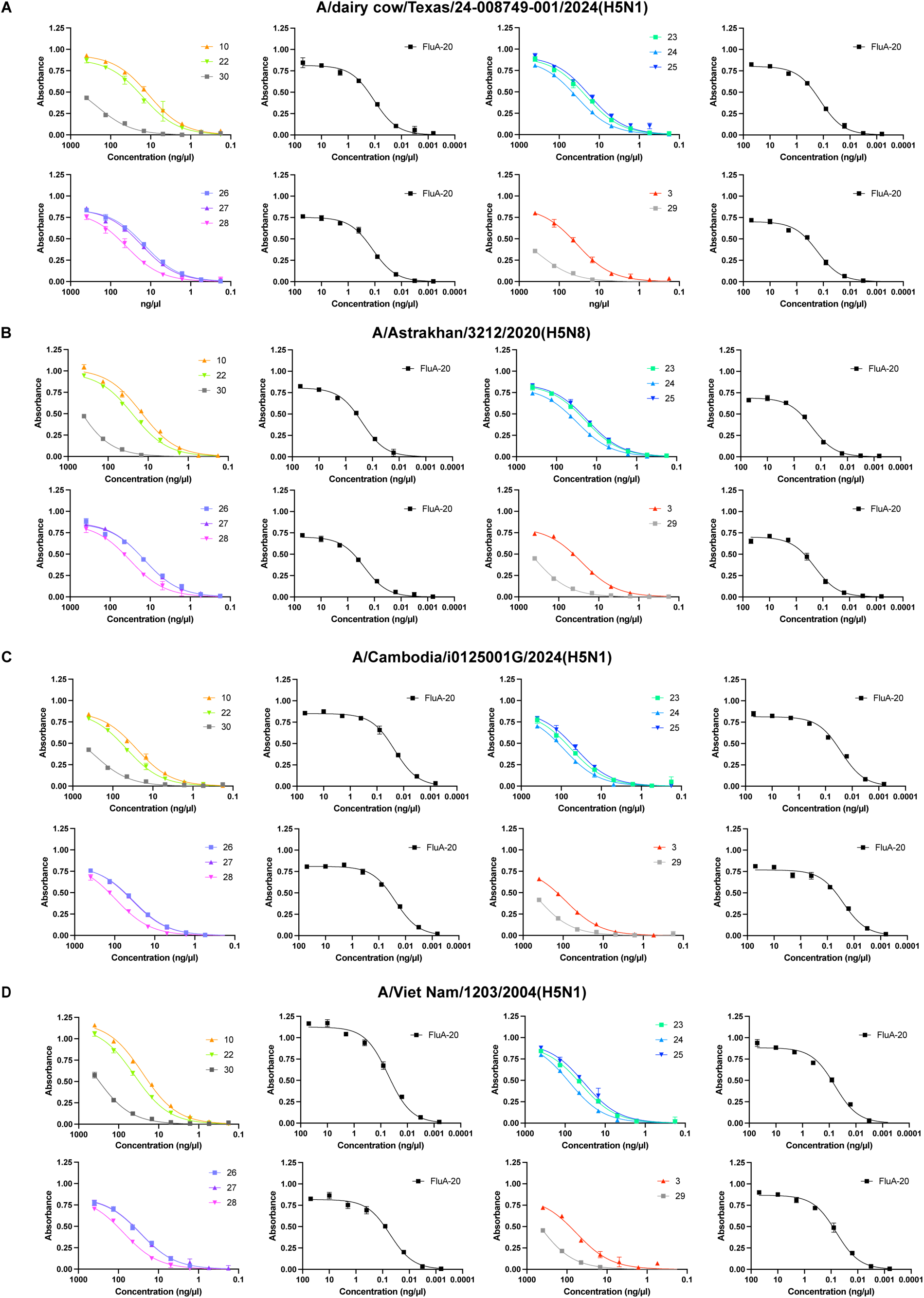
ELISA titration curves of purified IgG from milk samples to recombinant H5 HAs from the indicated isolates. (A) A/dairy cow/Texas/24-008749-001/2024(H5N1), (B) A/Astrakhan/3212/2020(H5N8), (C) A/Cambodia/i0125001G/2024(H5N1), (D) A/Viet Nam/1203/2004(H5N1). Data are background-subtracted absorbance values from three technical replicates. Curves for each ELISA plate are shown alongside standard curves for mAb FluA-20 (36) from the same plate.

**Figure S7.**
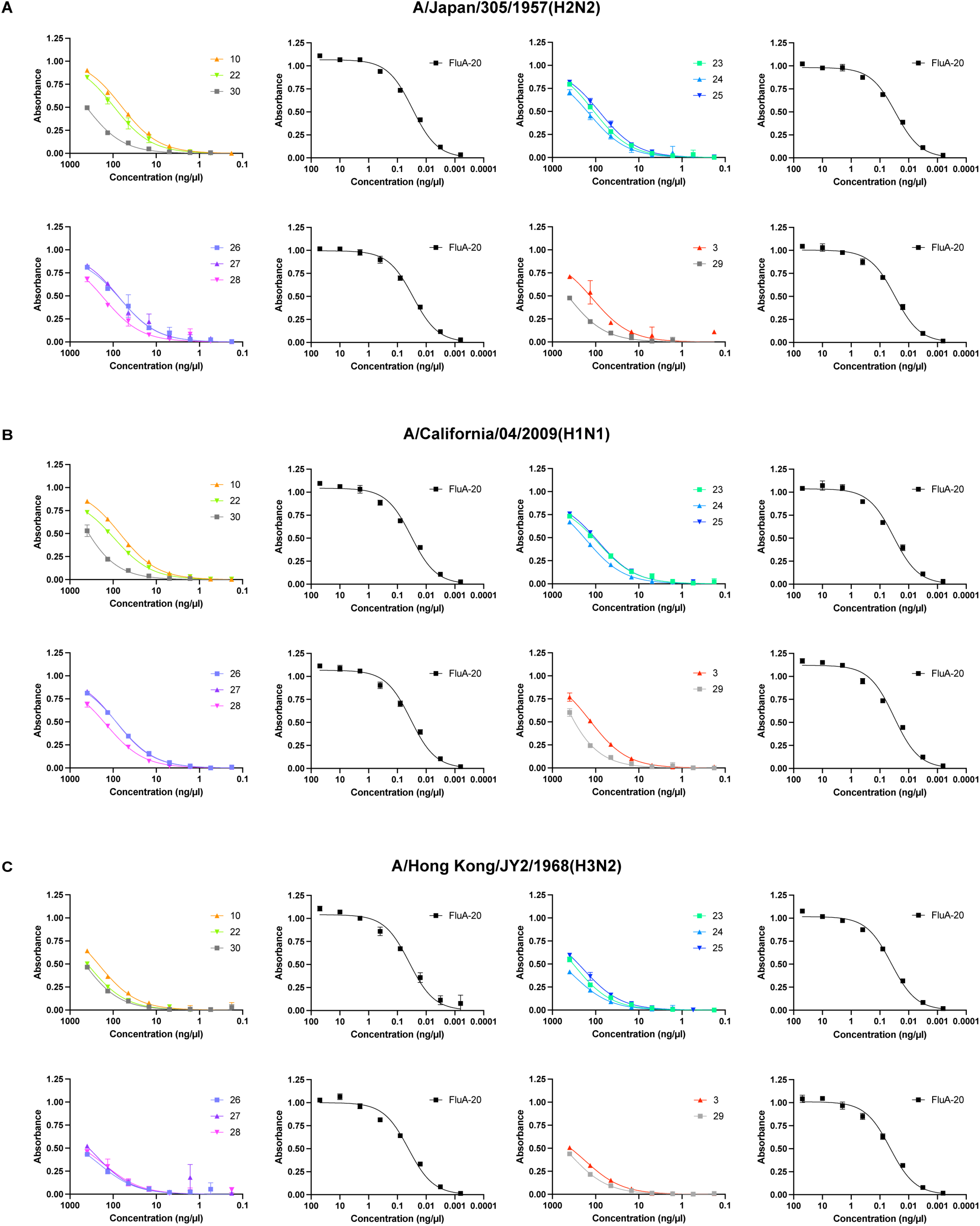
ELISA titration curves of purified IgG from milk samples to recombinant HAs from the indicated isolates. (A) A/Japan/305/1957(H2N2), (B) A/California/04/2009(H1N1), (C) A/Hong Kong/JY2/1968(H3N2). Data are background subtracted absorbance values from three technical replicates. Curves for each ELISA plate are shown alongside standard curves for mAb FluA-20 (36) from the same plate.

**Table S1.** Complete information for milk samples. “-”: information was unavailable. “*” Samples 6-8 are presumed to be from California, but brand and processing plant information were unavailable. Antibody positivity was determined by ELISA with A/dairy cow/Texas/24-008749-001/2024 H5 HA (See Figure S1). Brand and processing plants were deidentified and provided with numerical aliases.

| Sample Number | State | Brand alias | Plant alias | Milkfat | Pasteurization | Other processing | Expiration Date | Ab positivity (ELISA) |
| --- | --- | --- | --- | --- | --- | --- | --- | --- |
| 1 | PA | 1 | 1 | Whole | Pasteurized |  | 10/22/24 | Neg |
| 2 | CA | 2 | 2 | Skim | Pasteurized |  | 10/22/24 | Neg |
| 3 | CA | 3 | 2 | Skim | Pasteurized |  | 10/15/24 | Pos |
| 4 | CA | 4 | 3 | Whole | Pasteurized |  | 10/17/24 | Neg |
| 5 | CA | 5 | 4 | 2% | Ultra pasteurized | Lactose Free | 11/21/24 | Neg |
| 6 | CA* | - | - | Cream | - |  | - | Neg |
| 7 | CA* | - | - | Cream | - |  | - | Neg |
| 8 | CA* | - | - | 2% | - |  | - | Neg |
| 9 | CO | 6 | 5 | 2% | Ultra pasteurized |  | 3/7/25 | Neg |
| 10 | CO | 7 | 6 | Whole | Pasteurized |  | 10/21/24 | Pos |
| 11 | CO | 8 | 7 | Whole | Ultra pasteurized |  | 12/20/24 | Neg |
| 12 | CO | 9 | 7 | Whole | Ultra pasteurized |  | 12/2/24 | Neg |
| 13 | CO | 9 | 7 | 2% | Ultra pasteurized |  | 12/20/24 | Neg |
| 14 | CA | 10 | 8 | 2% | Ultra pasteurized |  | 12/10/24 | Neg |
| 15 | CA | 10 | 9 | Whole | Vat pasteurized |  | - | Neg |
| 16 | CA | 8 | 10 | Unk | - |  | - | Neg |
| 17 | CA | 11 | 11 | Whole | Pasteurized |  | 10/23/24 | Neg |
| 18 | CA | 8 | 3 | Whole | Pasteurized |  | 10/31/24 | Neg |
| 19 | MN | 8 | 12 | Whole | Pasteurized |  | 11/3/24 | Neg |
| 20 | MI | 12 | 13 | Whole | Ultra pasteurized | Lactose free, ultra filtered | 12/18/24 | Neg |
| 21 | OH | 13 | 14 | 2% | Ultra pasteurized |  | 10/29/24 | Neg |
| 22 | CO | 14 | 5 | Skim | Pasteurized |  | 11/23/24 | Pos |
| 23 | CO | 14 | 5 | Whole | Pasteurized |  | 11/28/24 | Pos |
| 24 | CO | 15 | 15 | 2% | Pasteurized |  | 11/20/24 | Pos |
| 25 | CO | 16 | 6 | 2% | Pasteurized |  | 11/18/24 | Pos |
| 26 | CO | 17 | 16 | Whole | Pasteurized |  | 11/22/24 | Pos |
| 27 | CO | 18 | 6 | 2% | - | Lactose Free | 11/10/24 | Pos |
| 28 | CO | 7 | 6 | Whole | Pasteurized |  | 11/20/24 | Pos |
| 29 | PA | 19 | 17 | Whole | Pasteurized |  | 11/27/24 | Neg |
| 30 | PA | 1 | 1 | Whole | Pasteurized |  | 11/29/24 | Neg |
| 31 | MI | 6 | 18 | Whole | Pasteurized |  | 12/5/24 | Neg |
| 32 | MI | 6 | 18 | 1% | Pasteurized |  | 12/4/24 | Neg |
| 33 | UT | 20 | 19 | Whole | Ultra pasteurized |  | 12/2/24 | Neg |
| 34 | MI | 12 | 18 | Whole | Ultra pasteurized | Lactose free, ultra filtered | 1/14/25 | Neg |
| 35 | KS | 3 | 20 | Whole | Ultra pasteurized |  | 1/17/25 | Neg |
| 36 | CO | 6 | 5 | Half & Half | Ultra pasteurized |  | 3/9/25 | Neg |

**Table S2.**
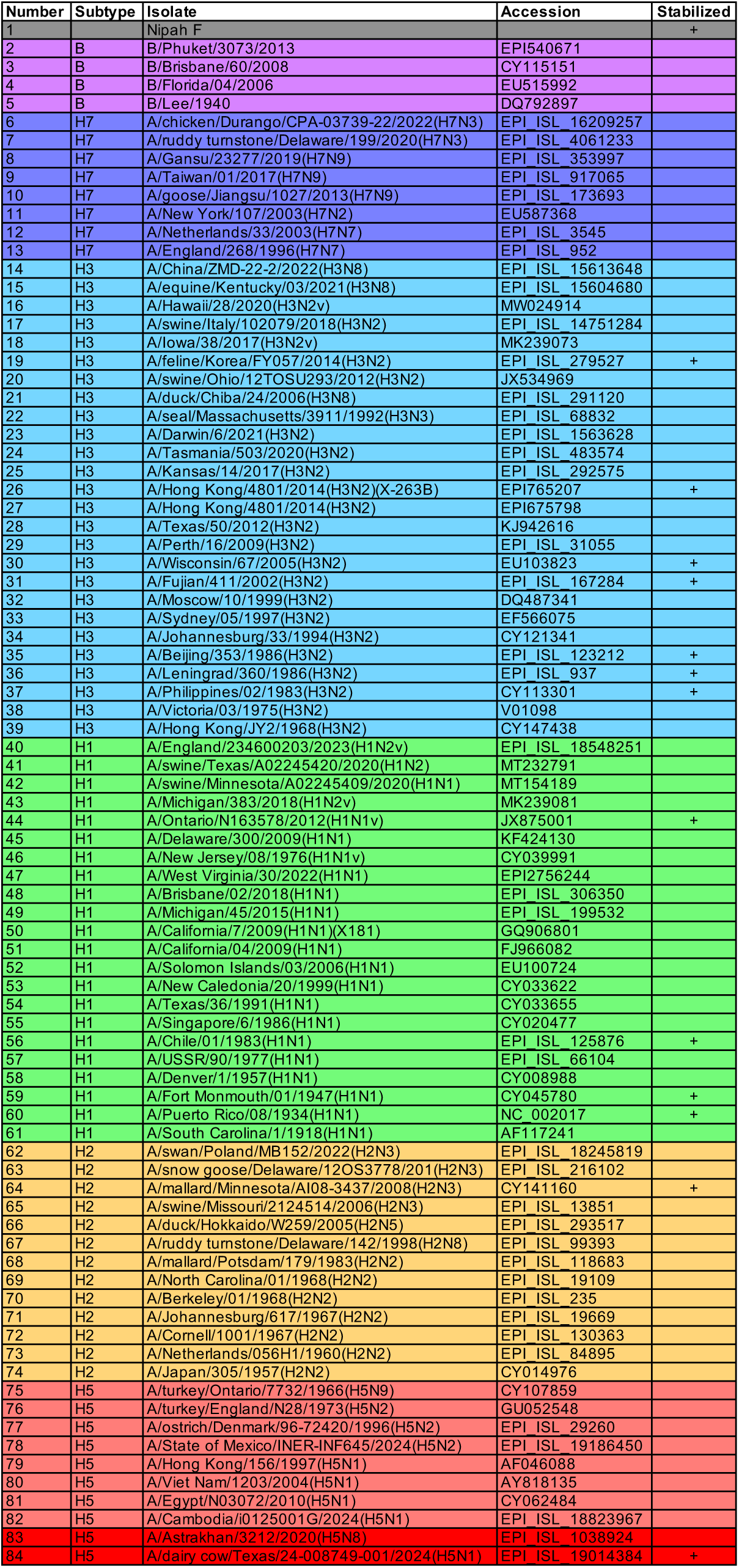
Information for recombinant HAs used for ELISAs. Nipah F was used a negative control. HAs indicated with a “+” contain stabilizing mutations (31, 32).

## References

1. T. Horimoto, Y. Kawaoka, Reverse genetics provides direct evidence for a correlation of hemagglutinin cleavability and virulence of an avian influenza A virus. J Virol 68, 3120–3128 (1994).

2. R. Xie et al., The episodic resurgence of highly pathogenic avian influenza H5 virus. Nature 622, 810–817 (2023).

3. USDA (2025) HPAI Confirmed Cases in Livestock.

4. L. C. Caserta et al., Spillover of highly pathogenic avian influenza H5N1 virus to dairy cattle. Nature 634, 669–676 (2024).

5. E. R. Burrough et al., Highly Pathogenic Avian Influenza A(H5N1) Clade 2.3.4.4b Virus Infection in Domestic Dairy Cattle and Cats, United States, 2024. Emerg Infect Dis 30, 1335-1343 (2024).

6. CDC (2025) H5 Bird Flu: Current Situation.

7. A. M. Mellis et al., Serologic Evidence of Recent Infection with Highly Pathogenic Avian Influenza A(H5) Virus Among Dairy Workers - Michigan and Colorado, June-August 2024. MMWR Morb Mortal Wkly Rep 73, 1004-1009 (2024).

8. I. Shittu et al., A One Health Investigation into H5N1 Avian Influenza Virus Epizootics on Two Dairy Farms. Clin Infect Dis 10.1093/cid/ciae576 (2024).

9. J. Leonard et al., Notes from the Field: Seroprevalence of Highly Pathogenic Avian Influenza A(H5) Virus Infections Among Bovine Veterinary Practitioners - United States, September 2024. MMWR Morb Mortal Wkly Rep 74, 50-52 (2025).

10. R. G. Webster, V. S. Hinshaw, W. J. Bean, G. Sriram, Influenza viruses: transmission between species. Philos Trans R Soc Lond B Biol Sci 288, 439–447 (1980).

11. C. A. Russell et al., Improving pandemic influenza risk assessment. Elife 3, e03883 (2014).

12. D. J. Smith et al., Mapping the antigenic and genetic evolution of influenza virus. Science 305, 371–376 (2004).

13. K. B. Westgeest et al., Genetic evolution of the neuraminidase of influenza A (H3N2) viruses from 1968 to 2009 and its correspondence to haemagglutinin evolution. J Gen Virol 93, 1996–2007 (2012).

14. M. R. Sandbulte et al., Discordant antigenic drift of neuraminidase and hemagglutinin in H1N1 and H3N2 influenza viruses. Proc Natl Acad Sci U S A 108, 20748–20753 (2011).

15. J. E. Salk, P. C. Suriano, Importance of antigenic composition of influenza virus vaccine in protecting against the natural disease; observations during the winter of 1947-1948. Am J Public Health Nations Health 39, 345–355 (1949).

16. A. C. Tricco et al., Comparing influenza vaccine efficacy against mismatched and matched strains: a systematic review and meta-analysis. BMC Med 11, 153 (2013).

17. F. Peña-Mosca et al., The impact of influenza A H5N1 virus infection in dairy cows. Research Square (2025).

18. N. J. Halwe et al., H5N1 clade 2.3.4.4b dynamics in experimentally infected calves and cows. Nature 637, 903–912 (2025).

19. A. L. Baker et al., Dairy cows inoculated with highly pathogenic avian influenza virus H5N1. Nature 637, 913–920 (2025).

20. J. U. Oguzie et al., Avian Influenza A(H5N1) Virus among Dairy Cattle, Texas, USA. Emerg Infect Dis 30, 1425–1429 (2024).

21. X. Xiong et al., Structures of complexes formed by H5 influenza hemagglutinin with a potent broadly neutralizing human monoclonal antibody. Proc Natl Acad Sci U S A 112, 9430–9435 (2015).

22. CFDA (2025) H5N1 Bird Flu Virus in Livestock. (California Department of Food and Agriculture).

23. L. R. Robinson-McCarthy, K. E. Zirckel, H. C. Simmons, V. Le Sage, K. R. McCarthy, A replicating recombinant vesicular stomatitis virus model for dairy cattle H5N1 influenza virus glycoprotein evolution. bioRxiv 10.1101/2025.02.27.640582, 2025.2002.2027.640582 (2025).

24. USDA (2025) The Occurrence of Another Highly Pathogenic Avian Influenza (HPAI) Spillover from Wild Birds into Dairy Cattle. (USDA Animal and Plant Health Inspection Service).

25. USDA (2025) APHIS Identifies Third HPAI Spillover in Dairy Cattle. (USDA Animal and Plant Health Inspection Service).

26. USDA (2025) APHIS Confirms D1.1 Genotype in Dairy Cattle in Nevada. (USDA Animal and Plant Health Inspection Service).

27. CEIRR (2024) Best Practice Guidelines for the Reporting of Milk Testing and Results. (Centers of Excellence for Influenza Research and Response).

28. U. J. Buchholz, S. Finke, K. K. Conzelmann, Generation of bovine respiratory syncytial virus (BRSV) from cDNA: BRSV NS2 is not essential for virus replication in tissue culture, and the human RSV leader region acts as a functional BRSV genome promoter. J Virol 73, 251–259 (1999).

29. A. G. Schmidt et al., Preconfiguration of the antigen-binding site during affinity maturation of a broadly neutralizing influenza virus antibody. Proc Natl Acad Sci U S A 110, 264–269 (2013).

30. K. R. McCarthy et al., A Prevalent Focused Human Antibody Response to the Influenza Virus Hemagglutinin Head Interface. mBio 12, e0114421 (2021).

31. F. J. Milder et al., Universal stabilization of the influenza hemagglutinin by structure-based redesign of the pH switch regions. Proc Natl Acad Sci U S A 119 (2022).

32. P. O. Byrne et al., Structural basis for antibody recognition of vulnerable epitopes on Nipah virus F protein. Nat Commun 14, 1494 (2023).

33. H. Hu et al., A human antibody recognizing a conserved epitope of H5 hemagglutinin broadly neutralizes highly pathogenic avian influenza H5N1 viruses. J Virol 86, 2978–2989 (2012).

34. C. Dreyfus et al., Highly conserved protective epitopes on influenza B viruses. Science 337, 1343–1348 (2012).

35. D. C. Ekiert et al., Antibody recognition of a highly conserved influenza virus epitope. Science 324, 246–251 (2009).

36. S. Bangaru et al., A Site of Vulnerability on the Influenza Virus Hemagglutinin Head Domain Trimer Interface. Cell 177, 1136–1152 e1118 (2019).

37. H. C. Simmons et al., A new class of antibodies that overcomes a steric barrier to cross-group neutralization of influenza viruses. PLoS Biol 21, e3002415 (2023).

38. J. ter Meulen et al., Human monoclonal antibody combination against SARS coronavirus: synergy and coverage of escape mutants. PLoS Med 3, e237 (2006).

